# Capturing instantaneous neural signal-behavior relationships with concurrent functional mixed models

**DOI:** 10.1101/2025.09.08.671383

**Authors:** Al W Xin, Erjia Cui, Francisco Pereira, Gabriel Loewinger

## Abstract

We previously proposed an analysis framework for fiber photometry data based on functional linear mixed models (FLMMs). Functional LMMs allow modeling associations between photometry traces and trial-specific scalar values like behavioral summaries and session number, while also accounting for between-animal heterogeneity. Here, we extend the method to concurrent FLMMs (cFLMMs), a method that can fit the instantaneous relationship between functional outcomes and functional covariates. Concurrent FLMMs enable testing of how the photometry signal is associated with, for example, a behavioral variable that evolves across within-trial timepoints (e.g. animal speed). cFLMMs can also model the relationship between the photometry signal and covariates in experiments with variable trial lengths (e.g., in studies where trials end when an animal responds). We illustrate the application of cFLMMs on two published studies and show the method can identify signal–behavior associations in analyses not possible with FLMMs. We find that analyzing photometry–behavior associations based on behavioral summaries (e.g., latency-to-response, average lick rate) can lead to misleading conclusions. We published our method in the `fastFMM` package, available as an R package through GitHub (https://github.com/awqx/fastFMM).

## Introduction

Fiber photometry allows for measurement of neural activity in vivo, and is useful for its high temporal resolution and neurotransmitter specificity. In ***Loewinger et al. (2025***), we demonstrated that *functional linear mixed models* (FLMMs) can effectively identify individual-level and trial-level temporal patterns in fiber photometry data. FLMMs provide key advantages over standard methods, including modeling instantaneous photometry signals at all the time points in each trial, providing hypothesis tests at each trial timepoint, and controlling for subject-specific effects. However, FLMMs are limited to capturing photometry dynamics with respect to scalar covariates. That is, FLMM can incorporate covariates that vary between trials, but they can only take a single value per trial, e.g., trial number, session, or behavioral summaries like latency-to-press, or total button presses during the trial. Limiting functional analysis to scalar covariates can coarsen available data and obscure time-varying associations between the signal and covariates.

In this work, we extend our method to *concurrent FLMMs* (cFLMMs), which can capture relationships with *functional covariates*, i.e. covariates that change across timepoints within one trial. Functional covariates can include (1) behavioral measures, like a subject’s speed, position, or actions, (2) changing experimental conditions, such as differing trial lengths or reward periods, or (3) an additional neural signal. Concurrent functional linear mixed modeling is one of a number of possible strategies for jointly modeling functional covariates and functional outcomes. For example, one could instead apply function-on-function regression (***Scheipl et al.,2016***), but we are not aware of longitudinal versions of this method. We opted for concurrent FLMMs because they can model instantaneous associations between the output and functional covariate (e.g., the relationship between the photometry signal and behavior at a specific within-trial timepoint) in longitudinal datasets.

### Box 1.

**Procedure for fitting a concurrent FLMM.**

We developed a procedure that fits a concurrent FLMM in three steps:

1. **Fit univariate models**. Separate linear mixed models are fit at every point along the *functional domain*. That is, if there are *L* longitudinal data points, *L* different linear mixed models will be produced.
2. **Smooth coefficients**. Aggregate and smooth the coefficient estimates for the linear models over the functional domain using penalized splines (***Cui et al. (2022***); ***Loewinger et al. (2025***)).
3. **Build joint confidence intervals**. Combine the within-timepoint variance and the between-timepoint covariance to create confidence bands around the smoothed estimates from Step 2.

The procedure for fitting a concurrent FLMM and a non-concurrent FLMM are similar and differ primarily in the calculation of joint confidence intervals in Step 3 (Concurrent FLMM estimation). In fact, if the functional covariate of interest is a scalar constant across the domain, the models fit by the concurrent and non-concurrent procedure are identical.

By introducing cFLMMs in the photometry context, we aim to demonstrate that they can capture temporal dynamics on real experimental data, similar to our experiments with *non-concurrent* FLMMs (ncFLMMs). Our examples are chosen to illustrate particular cases where concurrent modeling can provide insight into questions where modeling scalar covariates in ncFLMMs can produce misleading interpretations of coefficients. First, we will demonstrate how cFLMM captures the instantaneous relationship between the photometry signal and within-trial licking behavior. We contrast the cFLMM approach with ncFLMM, showing that the interpretation of ncFLMM coefficients can be biased depending on the choice of behavioral summary (i.e. average lick rate) (***Loewinger et al. (2025***), ***Jeong et al. (2022***)). Second, we will demonstrate how concurrent models can be deployed in experiments with variable trial length. We provide an example model for a foraging-like reward-seeking task where mice arrive at a reward zone at different trial timepoints (***Machen et al.,2025***) and evaluate the coefficient estimates against non-functional linear mixed model and ncFLMM baselines. Through these examples, we demonstrate that concurrent FLMMs can model a richer set of questions by allowing flexibility in controlling for changing conditions. This would be difficult to model with scalar covariates summarizing across within-trial timepoints.

## Results and discussion

### Concurrent FLMM modeling

Concurrent functional linear mixed modeling proceeds through the three broad steps of fitting univariate models, smoothing coefficients, and creating joint confidence intervals (Box 1), similar to the FLMM procedure described in ***Loewinger et al. (2025***). However, unlike our previous approach, concurrent models can take in experimental variables that vary across within-trial timepoints.

To describe this procedure, we can consider a hypothetical experiment investigating how the photometry signal changes based on a mouse’s instantaneous velocity during the experiment, illustrated in Figure 1A-i. In the cFLMM, we model the association of photometry signal at each time point with the instantaneous mouse velocity at the same time point. More formally, let *i* index a mouse ID, *j* index trial number, and *s* indicate a specific time during trial *j* (i.e., the functional domain). We model the instantaneous photometry outcome for mouse *i* on trial *j* at trial timepoint *s, Y*_*i,j*_(*s*), as a function of the instantaneous mouse velocity *X*_*i,j*_(*s*), as shown in Fig 1A-ii. The notation (*s*) in the mouse velocity variable *X*_*i,j*_(*s*) indicates that its value can evolve across within-trial time-points, like the photometry signal, *Y*_*i,j*_(*s*). Our model allows the photometry–speed association to evolve across trial timepoints through the functional intercept *β*_0_(*s*), and the functional slope *β*_1_(*s*). The subject-specific random intercept *γ*_*i*,0_(*s*) and subject-specific random slope *γ*_*i*,1_(*s*) allow for the magnitude of these associations to vary across animal *i* and timepoints *s*. Finally, we denote random noise as *ϵ*_*i,j*_(*s*), and formulate the model as:

**Figure 1.**
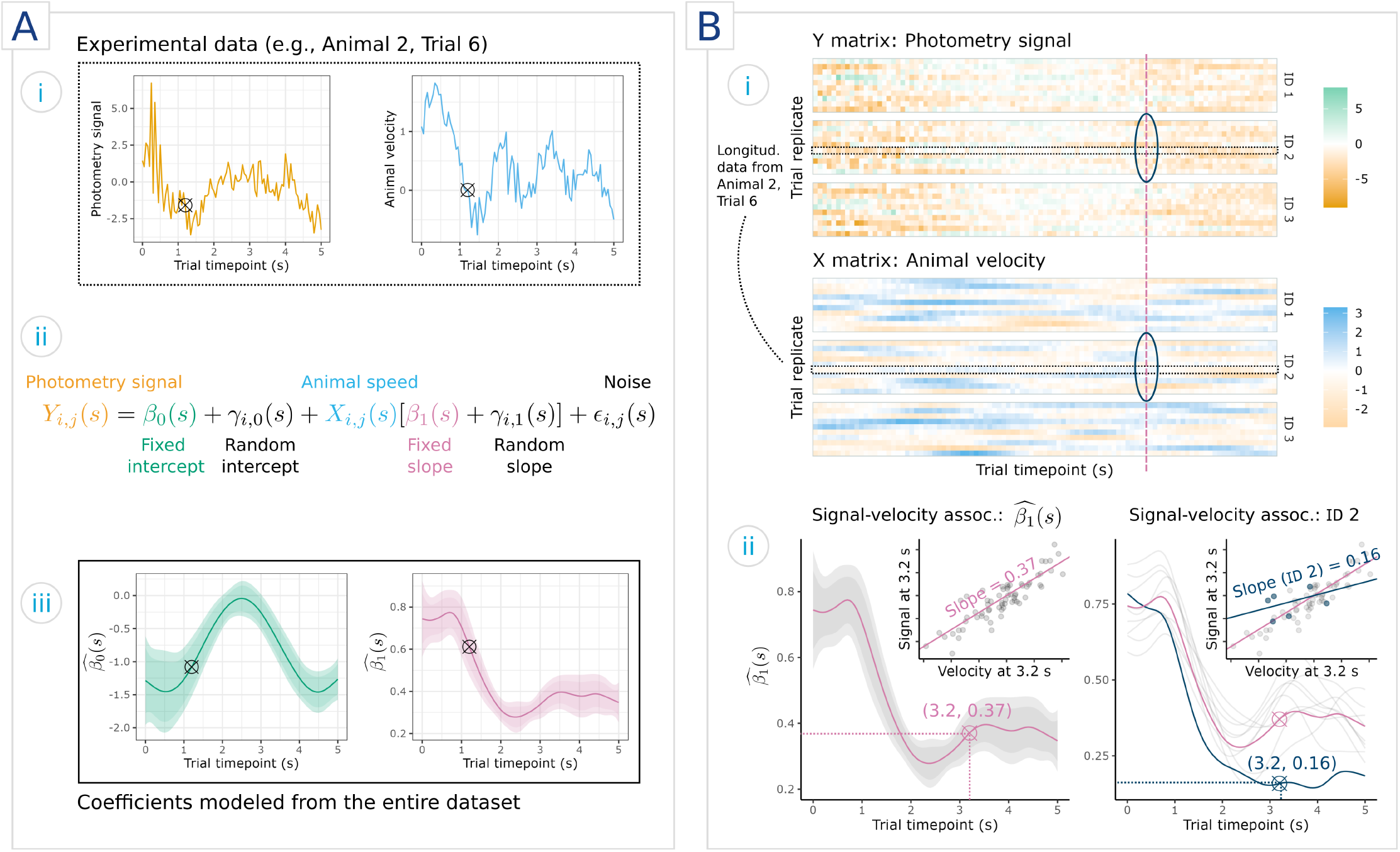
Visualization of an example concurrent functional linear mixed model on synthetic data. **A**) **A-i**) Functional outcomes (photometry signal) and inputs (animal velocity) over one trial, **A-ii**) labeled model equation for the time point marked with cross-hairs in (i) and (iii), **A-iii**) output functional fixed coefficients *β*_0_(*s*) and *β*_1_(*s*). **B**) Example experimental data and interpretation of effects, including the **B-i**) heat maps of the photometry dataset and speed dataset (groups are subjects *i*, rows are trials *j*, and columns are time points *s*), and **B-ii**) fixed effects at a given timepoint (left) and random effects at the same point (right).

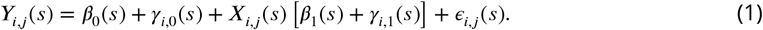

The outputs of this model are estimates 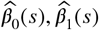 of the true fixed effect functional coefficient, *β*_0_(*s*) and *β*_1_(*s*), shown in Figure 1A-iii. We refer to these as *functional* regression coefficients because their values can vary over the course of trial timepoints, *s*. The estimated functional intercept 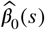 can be interpreted as the average signal at *s* when the velocity at *s* is zero (i.e., *X*_*i,j*_(*s*) = 0). The estimated functional slope 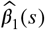 is the average change in signal associated with a one unit change in velocity at *s*. For example, in Figure 1A-iii, 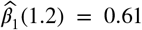, so a one-unit change in velocity is associated with a 0.61 unit increase in mean photometry signal at 1.2 seconds into the trial. This estimate is significantly higher than later in the trial (for example, 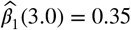), indicating that the association between signal and velocity becomes weaker as the trial progresses.

The model is fit to a dataset comprised of many trials for each of several animals, as shown in Figure 1B-i. While the fixed effects *β*_0_(*s*), *β*_1_(*s*) are pooled across all trials/animals, the random effects *γ*_*i*,0_(*s*), *γ*_*i*,1_(*s*) are perturbations to the fixed effects to represent animal-level differences, as illustrated in Figure 1B-ii. On the left plot of *β*_1_(*s*), the inset shows that 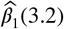 is the estimated slope between photometry signal and velocity at *s* = 3.2 seconds. Likewise, on the right plot of 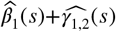, the inset shows how the random effect allows the slope to differ between animals. Each animal’s random effects are depicted in light gray, showing how animal-specific slopes are centered around the population fixed effect estimates. See Appendix 1 for more details on generating data for this demonstration.

### Modeling behaviors that change over the trial

To illustrate how concurrent models capture effects of behaviors, we reanalyzed data from a previously published study on mesolimbic dopamine dynamics in learning (***Jeong et al.,2022***). In this experiment, head-affixed mice were provided with sucrose solution at random times with 100 sucrose deliveries per session. The authors of ***Jeong et al. (2022***) were interested in modeling how the dopamine signal changes over the course of learning, i.e., modeling photometry signal as a function of session, trial, and inter-reward interval. Variable reward timing can complicate the analysis, as the signal is aligned to when the mouse licks the sucrose delivery mechanism rather than the moment when the sucrose is available. Furthermore, although mice were more prone to licking after reward delivery, each trial contained random unrewarded licking behavior before and immediately after delivery, as shown in Figure 2A-i. Previously, ***Jeong et al. (2022***) modeled the signal, shown in Figure 2A-ii, by averaging the photometry response in a predefined period after the first post-reward lick. In ***Loewinger et al. (2025***), we modeled the entire functional outcome of photometry. We aligned trials to the first lick, and estimated the association between the inter-reward interval and photometry signal. In contrast, here we centered the signal to reward delivery and then estimated the association between moment-by-moment lick behavior and the photometry signal.

**Figure 2.**
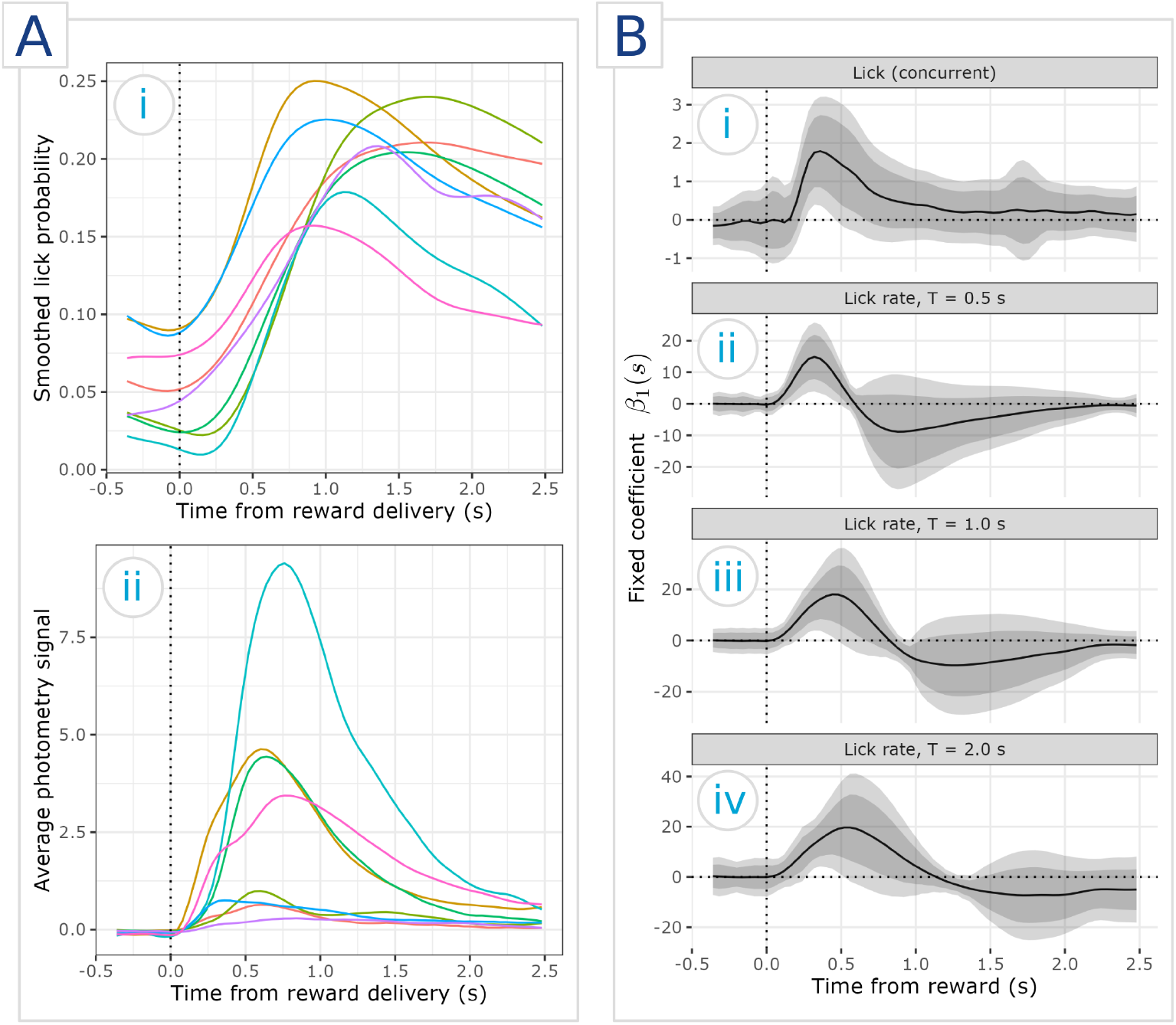
Fitted coefficients for concurrent and non-concurrent models of the expected signal with respect to licking behavior in mice randomly delivered sucrose solution. **A**) Features of the experimental data. **A-i**) The distribution of licking behaviors, averaged within mice and smoothed for readability, with one curve per mouse. **A-ii**) Photometry signal, averaged within mice. **B**) Estimates of the functional fixed coefficient *β*_1_(*s*) corresponding to different choices of FLMM and behavioral covariate. **B-i**) Concurrent model with an instantaneous functional covariate tracking licking behavior. **B-ii-iv**) Non-concurrent models with a trial-specific scalar covariate for the lick rate **ii**) 0.5 seconds after reward delivery, **iii**) 1.0 seconds after reward delivery, and **iv**) 2.0 seconds after reward delivery. Fixed effect estimates for trial and session can be found in Appendix 3.

#### Photometry signal as a function of intermittent licking behavior

To align with previous analyses in ***Loewinger et al. (2025***) and ***Jeong et al. (2022***)’s interest in longitudinal effects on photometry signal, we incorporated trial-specific covariates for trial number and session, modeling these as increasing numerical values rather than identical categorical variables. Modeling the experiment with a concurrent FLMM provides several advantages over the previous models. The time-dependence of the relationship between signal and licking behavior can be visualized without incorporating an additional covariate of reward-lick latency, which summarizes the trial-level lick timeseries. Unlike previous analyses, the concurrent model does not require discarding trials where the first lick occurs too quickly after the reward delivery. The concurrent model can directly estimate that the dopamine signal contribution of licking behavior (*β*_1_(*s*)) is zero prior to and immediately after reward delivery, rises 0.2 seconds after the reward is delivered, and peaks at around 0.3 seconds after reward delivery (Fig. 2B-i). We also include the fixed effects for trial and session in Appendix 3.

#### Comparison to non-concurrent models

We conducted a sensitivity analysis to assess how fitting a non-concurrent FLMM model would influence the coefficients and interpretation. We summarized licking behavior as a trial-specific scalar covariate, calculated as the average lick rate in a defined reward period *T* after sucrose delivery. We compared coefficient estimates for models of *T* = 0.5, 1.0, 2.0 seconds (Fig. 2Bii-iv).

While the trends of the fixed coefficients are similar to the concurrent model, the peak of the estimated effect is dependent on the choice of *L*. As reward periods are longer, the estimated peak occurs later. This trend may be due to different underlying lick dynamics resulting in the same summary statistic, leading to a flattening of coefficient estimates along the region used to summarize the behavioral data.

Furthermore,the point estimates of the ncFLMM fixed effect coefficients are negative later in the trial, unlike the concurrent model. However, because the underlying summary covariate analyzed in the ncFLMMs ignores these regions entirely, the modeled coefficients outside of the period *L* may be deceptive, as behavioral data in this region do not contribute to the average lick rate summary statistic. Although the point estimates imply that increased lick rates later in the trial are associated with lower expected signal outcome, it may be possible that the expected signal outcome is positive, but the lick rate is lower outside of *L*, causing the apparent negative effect. Although ncFLMM avoids the need to summarize photometry signal over the trial, the resulting models may still be biased due to transforming functional covariates into trial-specific scalars.

### Modeling behaviors in trials of varying lengths

In some experimental designs, trial length differs due to behavior or experimental conditions. Concurrent FLMMs can be applied to avoid issues with devising a fair summarization statistic of the photometry signal or behavior. To demonstrate this, we carried out an example cFLMM analysis in an experiment where mice receive rewards at inconsistent times. Mice were trained to wait in a trigger zone, run through a corridor, then receive a food reward in the reward delivery zone. Investigators were interested in testing whether the mean photometry signal changed based on the speed at which mice through the corridor, or the reward delivered (either a strawberry milkshake solution (SMS) or water. ***Machen et al. (2025***) tests more reward conditions, such as a non-caloric but sweet reward, though these are excluded for brevity. Within this setup, a natural way to define trials would be to summarize the signal over the period when the mouse is in the reward zone. However, this approach produces trials of different lengths, introducing bias depending on the choice of summary statistic.

#### Baseline 1: Non-functional linear mixed model

We demonstrated that summarizing functional data over time can be unreliable in this setup by producing two non-functional linear mixed models (LMMs) with different approaches to summarizing experimental variables (Eqn. 6). As before, let *s* indicate the trial timepoint, *i* index the animal ID, and *j* index the trial number. Let *S*_*i,j,R*_ be the set of timepoints where mouse *i* is in the reward zone in trial *j*. We compared models where the summary statistic was the average 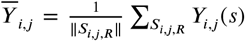 or the sum 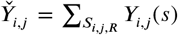 of photometry signals over the reward zone. For both choices, we found a strongly significant signal increase associated with the SMS reward (*p* < 0.0001). For 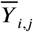, we found that the point estimates for latency and the interaction of the SMS reward and latency were both positive (Fig. 3A-i). Conversely, with the sum 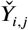, these point estimates were both negative (Fig. 3A-ii). This difference suggests that a sum over the period may overestimate the importance of arriving at the reward early, while an average signal penalizes an early arrival. In practice, choosing a summary may require prior knowledge of signal dynamics during the reward period, which may introduce bias in interpretation.

**Figure 3.**
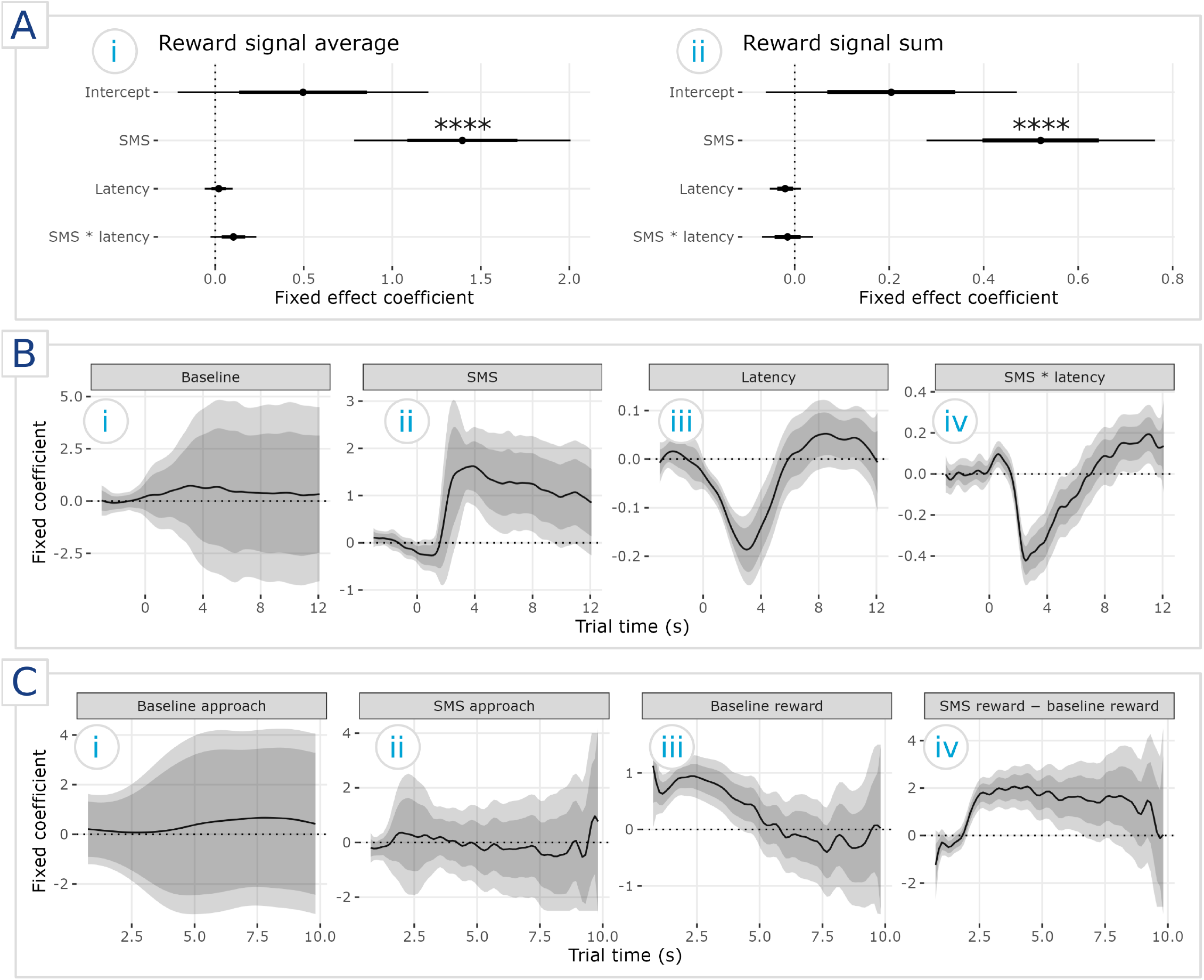
Comparisons of coefficient estimates for an experiment where mice receive different rewards at variable times. **A** Fixed coefficients in a non-functional linear mixed model fit to **A-i** the average of the photometry signal in the reward period or **A-ii** the sum, with *∗∗∗∗*: *p* ≤ 0.0001. **B** Functional fixed coefficients in a non-concurrent functional linear mixed model, corresponding to **B-i**) the intercept, i.e., the expected signal when latency is average and the mouse is given water; **B-ii**) the expected signal when latency is average and the mouse is given SMS; **B-iii**) the slope of mean signal change with respect to (w.r.t.) change in latency; and **B-iv**) the slope of mean signal change w.r.t. change in latency when the mouse is rewarded with SMS. C Functional fixed coefficients in a concurrent functional linear mixed model, corresponding to **C-i**) the intercept, i.e., the expected change in signal when a mouse in the baseline water condition has not yet received a reward; **C-ii**) unrewarded SMS, i.e., the expected change in signal when a mouse in the SMS condition has not yet received a reward; **C-iii**) baseline reward change, i.e., the expected difference in signal between a mouse receiving water and not receiving water; and **C-iv**) SMS reward change minus baseline reward change, i.e., the expected difference between receiving SMS and not receiving SMS and the value described in **A-iii**.

#### Baseline 2: Non-concurrent FLMM interaction model

Although a non-concurrent FLMM can capture the functional associations between the signal and behavioral data, the model produces coefficient estimates that could lead to misleading scientific conclusions. Because the non-concurrent model cannot use the moment-by-moment mouse location data, we summarized the behavior of each trial as a single scalar value of latency, i.e., how long it took for a mouse to reach the reward region from trial onset (Fig 3B) (Eqn. 7). Furthermore, for ease of interpretation, we centered trial latency. In the non-concurrent model, there are significant signal effects associated with the SMS reward, latency, and their interaction (Fig 3B). On trials with average latency, there is a significant increase in signal for mice receiving SMS (Fig 3B-ii) versus receiving the water baseline (Fig 3B-i) between 2 and 8 seconds into the trial. Increasing latency is associated with a negative effect on signal starting at trial start (zero seconds), reaching a minimum at around three seconds into the trial, and returning to zero at around 6 seconds into the trial (Fig 3B-iii). The shape of the fixed coefficient estimate for the interaction of SMS reward and latency is similar, though becomes slightly positive near the end of the trial (Fig 3B-iv).

#### cFLMM incorporating trial lengths as a functional interaction

For our concurrent model, we incorporated a binary functional covariate indicating whether the mouse had received the reward. We modeled an interaction between the reward indicator and the reward type (SMS or water) (Eqn. 8 in Variable trial length analysis). Unlike non-concurrent FLMMs, there is no expected difference in signal when a mouse has not yet received a reward, regardless of reward type (water or SMS) (Fig 3C-i, C-ii). However, expected signal change is positive early in the trial when mice receive a baseline reward (Fig 3C-iii), with an additional positive effect when the reward is SMS (Fig 3C-iv).

#### Interpreting non-concurrent FLMM vs. cFLMM coefficients

The concurrent model coefficients are more easily interpretable because the model separates the experiment into the relevant, discrete events. The first two fixed coefficients, for example, correspond to the expected signal for mice in a water trial that have not received a reward (Fig 3C-i) and mice in a SMS trial that have not received a reward (Fig 3C-ii). For both conditions, the estimated coefficients were not significantly different from zero and confirmed the experimental assumption that mice have no knowledge of what trial they are in before they receive a reward. The last two fixed coefficients, corresponding to signal for baseline reward (Fig 3C-iii) and the difference of SMS reward and baseline reward (Fig 3C-iv), separated the signal response attributable to receiving a reward in general versus receiving a desirable reward. Furthermore, the time dependence of all of these effects is captured by the functional nature of the coefficients and underlying covariates.

Within the non-concurrent FLMM, the instantaneous interpretation of the coefficients is less pertinent to the question of identifying the association between average signal and behavior. For example, the ncFLMM predicted no association when in a trial where water is the reward (Fig 3B-i) and a positive association in a trial where SMS is the reward (Fig 3B-ii). However, we know from analyzing the data that average latency across all mice and trials is also around 3 seconds. Thus, the observed peak in signal at around 3 seconds has an ambiguous interpretation, as it could be attributed to either receiving the SMS reward at that time or simply finishing the task.

Incorporating the estimated latency coefficients further complicates the interpretation of the ncFLMM. The dip in the estimated coefficients at around 3 seconds does not necessarily mean that mice with a slower-than-average latency will have a lower photometry response when they reach the reward. It could reflect that, in general, rewards increase the photometry signal magnitude and the slow mice have not yet reached the reward zone (Fig 3B-iii). The shape of the coefficient curve for the interaction `SMS * latency` also shows a dip, which seems to contradict the concurrent model’s estimate that SMS elicits a larger mean signal than water upon reward. However, the coefficients cannot be interpreted as effects associated with reward time, but only the effects at a specific trial time. The shape of the `SMS * latency` interaction functional coefficient can be explained as a correction for the strongly positive `SMS` main effect functional coefficient for slower-than-average mice. For example, a mouse receiving an SMS reward running a trial with a latency of 8 seconds does not exhibit, on average, a larger signal than on water trials at the trial timepoint of 4 seconds (doing so would imply prescience of the reward type). Thus, the negative `SMS * latency` coefficient estimate indicates that signal should be lower than what would be suggested by the `SMS` functional coefficient estimate at 4 seconds, as the `SMS` coefficient estimate reflects the estimated signal for mice running at an average latency.

#### Potential drawbacks of the functional covariate approach

For this particular problem, the non-concurrent model can model a larger functional domain than its concurrent counterpart. The functional coefficients for FLMMs are obtained by fitting individual linear mixed models at each point on the functional domain. When estimating the coefficients for a concurrent FLMM, each point must produce an identifiable model. This condition may be violated if a covariate has very low variance across every observation (e.g., mice never arrive at the reward zone early in the trial). Because all mice are approaching the reward at the start of trial and all mice are rewarded by the end of the trial, the concurrent FLMM cannot fit the extreme ends of the trial time frame. Here, this limitation is not relevant because there are likely no significant differences in mice when they are all experiencing the same condition. In this experiment in particular, mice are unaware of the reward they will receive when the experiment begins, and trials are unlikely to be meaningfully different within this time period.

The concurrent model also groups together all mice in the reward zone, which can suppress effects later in the trial. The indicator functional covariate is insensitive to how long ago a mouse has been reward, so a mouse that has just been rewarded and a mouse that has been rewarded much earlier in the trial are considered identical. Thus, the observed trend toward zero in the functional fixed effect estimates for the water reward and SMS reward may not indicate that mice have a lower photometry outcome response if they run slower (Figs. 3C-iii, -iv). Instead, the decrease may be partially attributed to more mice waiting in the reward period, diluting the observed signal outcome of newly-rewarded mice.

### Verification of method performance in simulations

Our concurrent FLMM approach produces both a *pointwise* and a *joint* confidence band for each coefficient time series. We produce the pointwise confidence band by estimating the variance of the fixed effects at each point and constructing appropriate 95% confidence intervals at each smoothed fixed effect estimate. The joint confidence band accounts for autocorrelation of signal values and produces a confidence band that corresponds to a hypothesis test across all timepoints simultaneously. To evaluate both the pointwise and joint confidence interval coverage of the concurrent FLMM models, we simulated data from a known model in a variety of realistic settings of dataset size and noise. We calculated pointwise coverage by finding the empirical coverage probability at each point and then averaging across the domain. In contrast, we assessed joint coverage by finding the empirical coverage probability of the confidence band, i.e., the probability that the entirety of the true fixed effects function lies within the estimated joint confidence band.

We simulated data generated from the following model

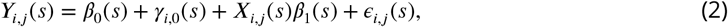

which contains a fixed intercept *β*_0_(*s*), fixed slope *β*_1_(*s*), and random intercept *γ*_0_(*s*). Within these simulations, we assumed that noise can arise from between-animal heterogeneity, e.g., individual differences affecting the magnitude of association between velocity and photometry signal. Furthermore, we also incorporated random measurement noise uncorrelated with experimental variables. These noise components were defined as 1) the signal-noise ratio of fixed effects to random effects `SNR`_*B*_, i.e., effect heterogeneity that arises from animal-level differences, and 2) the signal-noise ratio of true underlying values to random Gaussian noise `SNR`_*σ*_, i.e., the *ϵ*_*i,j*_ (*s*) term in Equation 2. We also refer to these components as measurement noise and between subject variability, respectively. We performed 200 replicates of 5, 10, …, 45, 50 subjects (simulated animals) with 100 trials each. Specific implementation details and examples of noisy data are included in Appendix 2.

We found that coverage generally asymptotes at ≥ 10 subjects for all noise conditions (Fig 4), though the value of the asymptotic coverage is dependent on the specific levels of noise. The signal-noise ratio of fixed effects to random effects (`SNR`_*B*_) shows a more pronounced effect on coverage than the relative importance of Gaussian noise (`SNR`_*σ*_) for both joint and pointwise coverage. Joint coverage tends to be overconfident when `SNR`_*B*_ = 1, close to theoretical 95% coverage when `SNR`_*B*_ = 1/2, and conservative when `SNR`_*B*_ = 1/3 and Gaussian noise is low (Fig 4A). Pointwise coverage tends to be conservative in all tested conditions except for `SNR`_*B*_ = 1 (Fig 4B).

**Figure 4.**
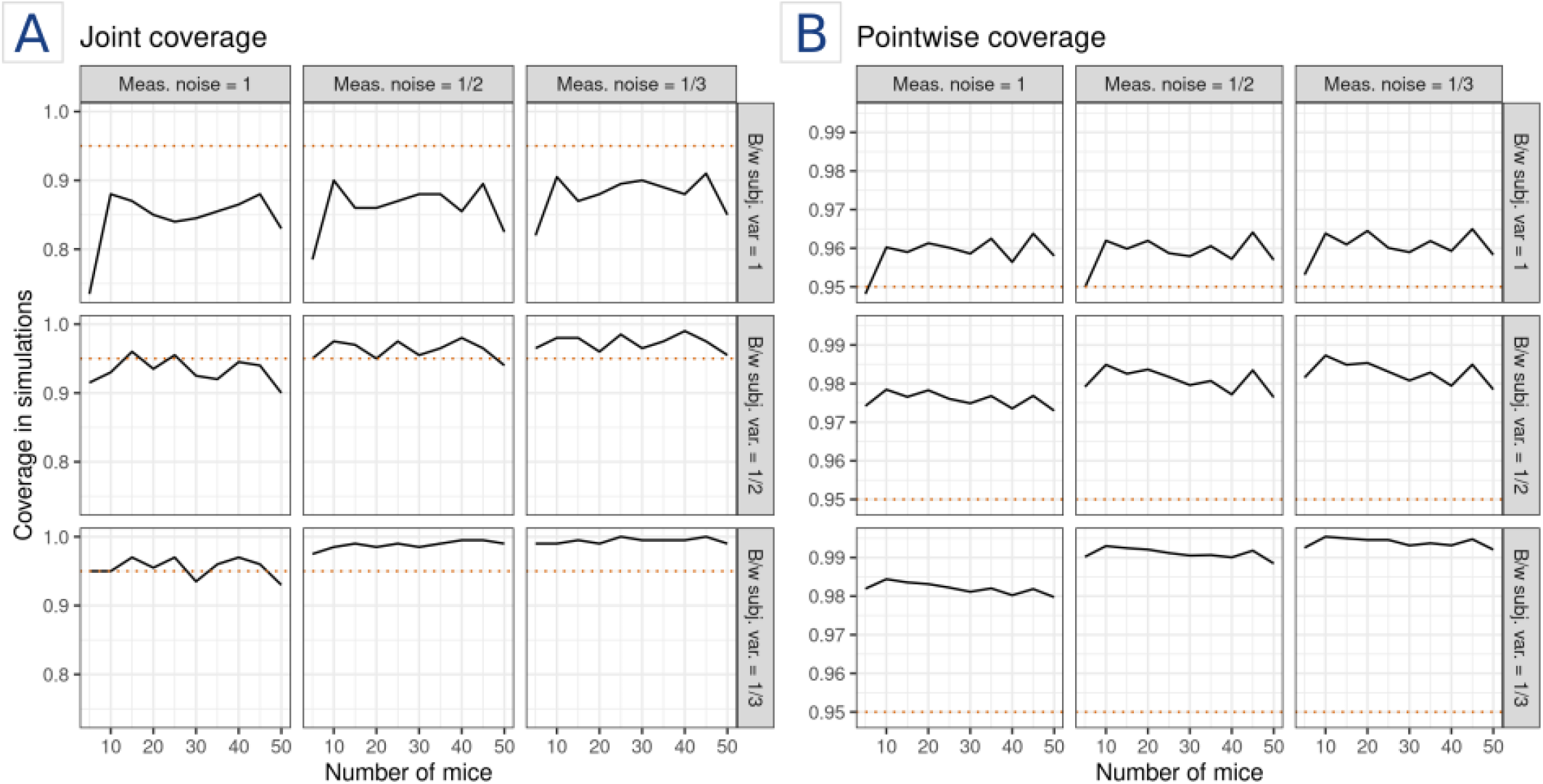
Nominal coverage achieved in simulations, reported as **A**) joint coverage and **B**) pointwise coverage for different values of measurement noise (meas. noise) and between subject variability (b/w subj. var.). Simulations were performed on 200 replicates of each condition. The horizontal red dotted line indicates the theoretical 95% coverage mark.

The simulation results suggest that relatively high levels of subject-specific effects impede coverage more than high levels of random measurement noise. Even in settings where the standard deviation of Gaussian noise is equal to that of the underlying signal (`SNR`_*σ*_ = 1, leftmost column in Fig 4), joint coverage achieves 95% for 15 subjects when `SNR`_*B*_ = 1/2 and 5 subjects when `SNR`_*B*_ = 1/3. However, when `SNR`_*B*_ = 1 (topmost rows in A, B of Fig 4), joint coverage peaks at around 90% regardless of the number of subjects or level of `SNR`_*σ*_. Barring high levels of between-subject variability, coverage is likely valid, albeit conservative.

## Materials and methods

### Concurrent FLMM estimation

#### General form of cFLMMs

Let *p* be the number of fixed effects, *q* be the number of random effects, *i* be the animal ID, *j* be the trial number, and *l* be the session number. We describe the concurrent functional linear mixed model of *Y*_*i,j,l*_(*s*), or the photometry signal at timepoint *s* for animal *i*, trial *j*, and session *l*), with the generic mean model

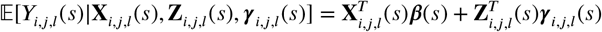

where subscripts reflect whether variables are subject-, trial-, or session-specific. In this formulation, **X**_*i,j,l*_(*s*) is the *p*×1 column vector of covariates for fixed effects, **Z**_*i,j,l*_(*s*) is the *q* ×1 column vector of covariates for random effects, ***γ***_*i,j,l*_(*s*) is the *q* × 1 column vector of random effects, and **β**(*s*) is the *p* × 1 column vector of fixed effects. We assume a Gaussian distribution for the conditional likelihood *Y*_*i,j,l*_(*s*)|**X**_*i,j,l*_(*s*), **Z**_*i,j,l*_(*s*), ***γ***_*i,j,l*_(*s*).

Throughout this paper, we write models as expressions with the explicit form

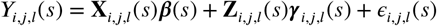

where we assume random effects ***γ*** _*i*_(*s*) ∈ ℝ ^*q*^ and noise terms ***ϵ*** (*s*) ∈ ℝ ^1^ are mutually independent and Normally distributed. Estimates for random effects **β** (*s*) and fixed effects ***γ***_*i*_ (*s*) are correlated across the functional domain. When describing cFLMM models, we replace the general vector notation **X**_*i,j,l*_ with individual covariate terms and descriptive names (e.g., `Lick`_*i,j,l*_(*s*)).

#### Methodological contributions to joint confidence interval estimation

We provide a high-level overview of the methodological details of the concurrent FLMM here and include a more detailed description in *Appendix 3* of ***Loewinger et al. (2025***). The main methodological contribution required for this approach is an extension of a method of moments covariance estimator needed to compute 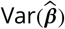 and calculate 95% confidence intervals. Briefly, after fitting pointwise linear mixed models, we obtain estimates of the fixed effects 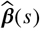 at each within-trial timepoint *s*. As in ***Loewinger et al. (2025***); ***Cui et al. (2022***), the covariance between fixed effects estimates can be described as

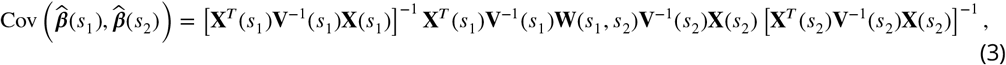

where we define

- **X**(*s*): the design matrix for the fixed coefficients at point *s*,
- **Z**(*s*): the design matrix for the random coefficients at point *s*,
- **V**(*s*) = **Z**(*s*)**H**(*s*)**Z**(*s*)^*T*^ + **R**(*s*), where **H**(*s*) and **R**(*s*) are the covariance matrices of ***γ***(*s*) and ***ϵ***(*s*), respectively and estimates 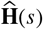 and 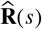 can both be obtained from the first step,
- **W**(*s*_1_, *s*_2_) = **Z**(*s*)**G**(*s*_1_, *s*_2_)**Z**^*T*^ (*s*), where **G** is the covariance matrix of random effects and **G**(*s*_1_, *s*_2_) = Cov(***γ***(*s*_1_), ***γ***(*s*_2_)), where ***γ***(*s*) is a one-dimensional vector concatenating the random effects for all subjects at point *s*.

In contrast with non-concurrent FLMMs, covariance estimation for cFLMMs incorporates unique **Z**(*s*) and **X**(*s*) values at each point on the functional domain. The number of calculations required to obtain an estimate grows combinatorially with respect to the size of the functional domain *L*, the number of random effects, and the number of fixed effects, resulting in a highly memory-intensive procedure. This step can become computationally infeasible for even moderately-sized datasets if carried out naively. A major methodological contribution of this work was deriving a computationally efficient procedure, described in Appendix 4.

#### R package for cFLMM fitting

We incorporated cFLMM fitting into our existing package `fastFMM`, published on GitHub `awqx/fastFMM`. Concurrent models can be fit with the same notation as ncFLMMs by adding the argument `concurrent = TRUE` to the function `fastFMM::fui`. The following sections will provide examples of the code needed to fit both ncFLMMs and cFLMMs with `fastFMM`.

### Reanalysis of photometry measures of dopaminergic neurons

When applying concurrent FLMMs to time-varying behaviors, we re-analyzed data from Experiment 3 published in ***Jeong et al. (2022***), as described in Modeling behaviors that change over the trial. We briefly explain the experimental design before describing the model that we applied. The original authors of ***Jeong et al. (2022***) presented head-affixed mice with sucrose solution, with delivery times distributed exponentially with a mean of 12 seconds and 100 sucrose rewards within a single session. In our analysis, we defined trials as a window 0.4 seconds before and 2.5 seconds after reward delivery, with overlapping trials discarded. We specified the concurrent model as:

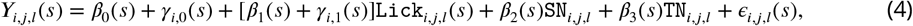

where *i* indexes the mouse ID, *j* indexes the trial number, and *l* indexes the session number. *Y*_*i,j,l*_(*s*) is the observed signal and *ϵ*_*i,j,l*_(*s*) is the observed noise at *s* for subject *i*, trial *j*, and session *l*. We included the covariates

- `Lick`_*i,j,l*_(*s*): the functional covariate of licking behavior, where `Lick`(*s*) = 1 when mouse *i* on trial *j* of session *l* is licking at time *s* and `Lick`(*s*) = 0 otherwise,
- `SN`_*i,j,l*_: the shifted and scaled session number, and
- `TN`_*i,j,l*_ the shifted and scaled trial number.

Regressors `SN, TN` are included because there may be temporal effects within a session (e.g. satiation), or between sessions (e.g. practice). We transformed `SN, TN`, by subtracting one and dividing by the maximum session number and trial number, respectively. This modification aids interpretation by setting the first session at zero and the max session as one.^1^

We included the random effects

- *γ*_*i*,0_(*s*): the random intercept for mouse *i* at time *s*. This is interpreted as the average photometry signal at *s* of mouse *i* on trials when when `Lick`_*i,j,l*_(*s*) = `SN`_*i,j,l*_ = `TN`_*i,j,l*_ = 0, and
- *γ*_*i*,1_(*s*) the random slope associated with the licking behavior for mouse *i* at time *s*, and the fixed coefficients

- *β*_0_(*s*): the intercept when `Lick`_*i,j,l*_(*s*) = `SN`_*i,j,l*_ = `TN`_*i,j,l*_ = 0, i.e., the mean signal at *s* on the first trial and first session when the mouse is not licking,
- *β*_1_(*s*): the mean effect on the signal associated with licking behavior at *s*,
- *β*_2_(*s*): the mean effect on the signal associated with the session number `SN`_*i,j,l*_ at time *s*, and
- *β*_3_(*s*): the mean effect on the signal associated with the trial number `TN`_*i,j,l*_ at time *s*.

We fit the concurrent model with the R code

~~~
fastFMM::fui(
formula = photometry ∼ session + trial + lick + (1 + lick | id),
data = data,
concurrent = TRUE
)
~~~

We fit the non-concurrent model

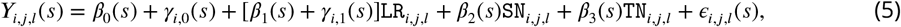

where we replaced `Lick`_*i,j,l*_ (*s*) with the scalar 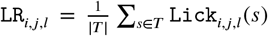, corresponding to the lick rate in the time region *T* after the reward delivery. To calculate |*T*|, we multiplied the number of seconds in *T* (either 0.5, 1.0, or 2.0) by the measurement frequency of 25 Hz. We fit the non-concurrent model with the R code

~~~
fastFMM::fui(
formula = photometry ∼ session + trial + LR + (1 + LR | id),
data = data,
concurrent = FALSE
)
~~~

where `LR` varied depending on the specified time period *T*.

### Variable trial length analysis

In Modeling behaviors in trials of varying lengths, we reanalyzed data from a foraging-like reward-seeking task, where mice run through a corridor and enter a reward zone at different times (***Machen et al.,2025***). Throughout this section, let *i* index the mouse and *j* index the trial. The non-functional linear mixed model contains no functional terms, and we defined it as

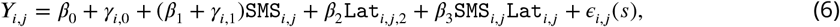

where the trial-specific scalar outcome *Y*_*i,j*_ is either 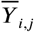 or 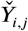, which denotes the average or the sum of the signal over the reward period, respectively. We defined the covariates

- `SMS`_*i,j*_: the trial outcome, where `SMS`_*i,j*_ = 1 for a trial with strawberry milkshake (SMS) and `SMS`_*i,j*_ = 0 for water, and
- `Lat`_*i,j*_: the latency, in seconds, until mouse *i* in trial *j* received the food reward.

For ease of interpretation, we centered the latency values, so `Lat`_*i,j*,2_ is the difference between the latency in that trial and the average latency across all trials. For example, `Lat`_*i,j*_ < 0 when mouse *i* in trial *j* ran the corridor faster than average and `Lat`_*i,j*_ > 0 when mouse *i* in trial *j* ran slower than average. The random effects are

- *γ*_*i*,0_: the random intercept for mouse *i*, interpreted as the average photometry signal of mouse *i* on trials with water reward and average latency, and
- *γ*_*i*,1_ the random slope for mouse *i*, interpreted as the subject-specific association between signal and the SMS reward for mouse *i* on average latency,

and the fixed effect coefficients are

- *β*_0_: the intercept when `SMS`_*i,j*_ = `Lat`_*i,j*_ = 0, i.e., the mean signal on water trials when a mouse responds with an average latency,
- *β*_1_: the mean difference in signals between SMS and water trials when a mouse responds with an average latency,
- *β*_2_: the change in mean signal associated with a one second increase in latency on water trials, and
- *β*_3_: the difference *A* − *B* between A) the change in mean signal associated with a one second increase in latency on SMS trials and B) *β*_2_.

We fit the two linear mixed models with the R package `nlme` (***Pinheiro et al.,2025***) with the code

~~~
lme_avg <- nlme::lme(
fixed = photo_avg ∼ SMS * latency,
random = ∼ outcome | id,
data = data
)
lme_sum <- nlme::lme(
fixed = photo_sum ∼ SMS * latency,
random = ∼ SMS | id,
data = dat
)
~~~

where the interaction term implicitly includes the individual terms `SMS` and `latency` in the model. Our non-concurrent FLMM treated the entire trial photometry signal as a functional outcome *Y*_*i,j*_(*s*). We specified the ncFLMM model as

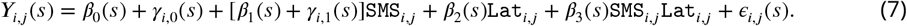

The interpretations of the random and fixed effects are analogous to the non-functional linear mixed model. However, in the functional setting, the coefficients correspond to associations between the signal at time *s* and the covariates. We fit the ncFLMM with the code

~~~
fastFMM::fui(
photometry ∼ SMS * latency + (SMS | id),
data = data,
concurrent = FALSE
)
~~~

where the interaction term implicitly includes the individual terms `SMS` and `latency` in the model.

The cFLMM model incorporated the functional covariate of `RZ`_*i,j*_ (*s*) (reward zone) instead of the scalar covariate `Lat`_*i,j*_. We specified the cFLMM model as

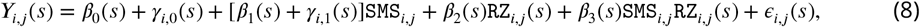

where `RZ`_*i,j*_ (*s*) = 0 if mouse *i* in trial *j* was in the corridor approaching the reward zone at time *s*, and `RZ`_*i,j*_(*s*) = 1 if mouse was in the reward zone at time *s*. The random effects are interpreted as

- *γ*_*i*,0_(*s*): the random intercept for mouse *i*, or the subject-specific average perturbation of *β*_0_(*s*) such that *β*_0_(*s*) + *γ*_*i*,0_(*s*) is the average signal for mouse *i* on water trials at time *s* when the reward has not yet been received, and
- *γ*_*i*,1_(*s*): the random slope for mouse *i*, or the subject-specific average perturbation of *β*_1_(*s*) such that *β*_1_(*s*) + *γ*_*i*,1_(*s*) is the average photometry signal change associated with being in an SMS trial for mouse *i* at time *s* when the reward has not yet been received,

and the functional fixed coefficients are interpreted as

- *β*_0_(*s*): the mean signal on a water trial at a timepoint *s* when the reward has not yet been received, i.e., when `RZ`_*i,j*_(*s*) = `SMS`_*i,j*_ = 0,
- *β*_1_(*s*): the mean difference in signal between SMS and water trials at a timepoint *s* when the reward has not yet been received,
- *β*_2_(*s*): the average difference between approach and reward in water trials at time *s*, and
- *β*_3_(*s*): the difference in differences *A* − *B* between A) the average signal difference between approach and reward on SMS trials, and B) the average difference in signal values between approach and reward on water trials (i.e., *β*_2_), at trial timepoint *s*.

Here, *β*_3_(*s*) = 0 when the association between reward type and the photometry signal does not change depending on the type of reward given. Conversely, *β*_3_(*s*) > 0 if, compared to water trials, SMS trials are linked with a greater change in mean signal between being in the reward zone versus the corridor at *s*. That is, *β*_3_(*s*) captures how the photometry signal is changed by a dynamic interaction between the type of reward and the behavior of the mouse. We fit the cFLMM with the code

~~~
fastFMM::fui(
photometry ∼ SMS * RZ + (SMS | id),
data = data,
concurrent = TRUE
)
~~~

where the interaction term implicitly includes the individual main effect terms `SMS` and `RZ` in the model.

## Acknowledgments

First, we thank Dr. Sofia Beas, the corresponding author and principal investigator of “The encoding of interoceptive-based predictions by the paraventricular nucleus of the thalamus D2R+ neurons”, for sharing experimental data and helping us interpret the results. We also thank Drs. Vijay Namboodiri and Huijeong Jeong, the corresponding authors of “Mesolimbic dopamine release conveys causal associations,” who generously shared their data and provided support for our previous publication, “A statistical framework for analysis of trial-level temporal dynamics in fiber photometry experiments”.

Simulation results in this study utilized the computational resources of the NIH HPC Biowulf cluster (https://hpc.nih.gov). This research was supported in part by the Intramural Research Program of the National Institutes of Health (NIH) (ZIC MH002968). The contributions of the NIH author(s) were made as part of their official duties as NIH federal employees, are in compliance with agency policy requirements, and are considered Works of the United States Government. However, the findings and conclusions presented in this paper are those of the authors and do not necessarily reflect the views of the NIH or the U.S. Department of Health and Human Services.

## Appendix 1

### Synthetic demonstration data

To construct synthetic data, we adapted the procedure described in ***Cui et al. (2022***) for concurrent modeling. Let *i* = 1, 2, …, *I* index subject IDs and *j* = 1, 2, …, *J* index replicates (i.e. trials). We simulated data according to the following functional mixed model

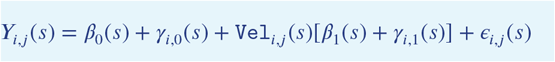

where we defined the components

- *Y*_*i,j*_(*s*): the observed photometry signal for animal *i* at trial *j* and time *s*;
- Vel_*i,j*_(*s*): the observed velocity for animal *i* at trial *j* and time *s*;
- *β*_0_(*s*): the functional intercept at time *s* across all trials and subjects;
- *β*_1_(*s*): the average change in mean signal associated with a one-unit increase in velocity at time *s* across all trials and subjects;
- *γ*_*i*,0_(*s*): the random functional intercept of mouse *i* at time *s* across all trials; and
- *γ*_*i*,1_(*s*): the random functional slope for mouse *i* at time *s*, i.e., the subject-specific perturbation to the population signal–velocity slope.

Let 𝒮 be the functional domain, with size *L*. Velocity values were drawn from 0.00 to 5.00 seconds in increments of 0.05 seconds, i.e., 𝒮 = {0.00, 0.05, …, 4.95, 5.00}, |𝒮| = *L* = 101. We sampled the vector of velocities across within-trial timepoints `Vel`_*i,j*_ = [`Vel`_*i,j*_(*s*_1_), …, `Vel`_*i,j*_(*s*_*L*_)]^*T*^ from the *L*-dimensional multivariate normal

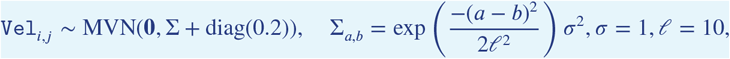

where we drew each vector `Vel`_*i,j*_ independently and identically distributed (i.i.d.) across values of *i* and *j*. We defined fixed effect as

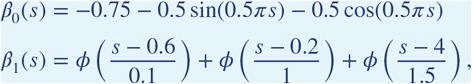

where *ϕ* is the standard Normal density function. We modeled random functional intercepts (*γ*_0,*i*_(*s*)) and slopes (*γ*_1,*i*_(*s*)) as the sum of two scaled orthonormal functions *ψ*_1_(*s*), *ψ*_2_(*s*). For every mouse *i* ∈ {1, …, *I*},

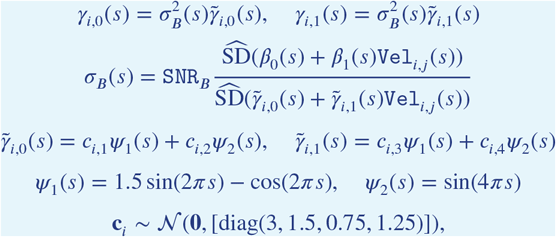

where 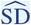 is the empirical standard deviation and `SNR`_*B*_ is a scaling parameter controlling the relative importance of fixed effects to subject-specific variability. Here, `SNR`_*B*_ = 1. We sampled the vectors **c**_*i*_ i.i.d. across subjects, *i*. We defined measurement noise *ϵ*_*i,j*_(*s*) as

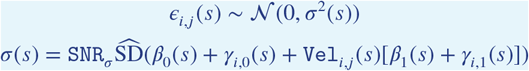

and sampled *ϵ*_*i,j*_(*s*) independently across mouse *i*, trial *j*, and time *s*. The scaling parameter `SNR`_*σ*_ controls the relative importance of the combined fixed and random effects to the measurement noise. Here, `SNR`_*σ*_ = 1. Unlike simulations in ***Cui et al. (2022***), 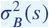 and *σ*^2^(*s*) are functions, allowing for `SNR`_*B*_ and `SNR`_*σ*_ to be constant along the functional domain. Visualizations of the random and fixed effects are below, in Figure 1.

**Appendix 1–figure 1.**
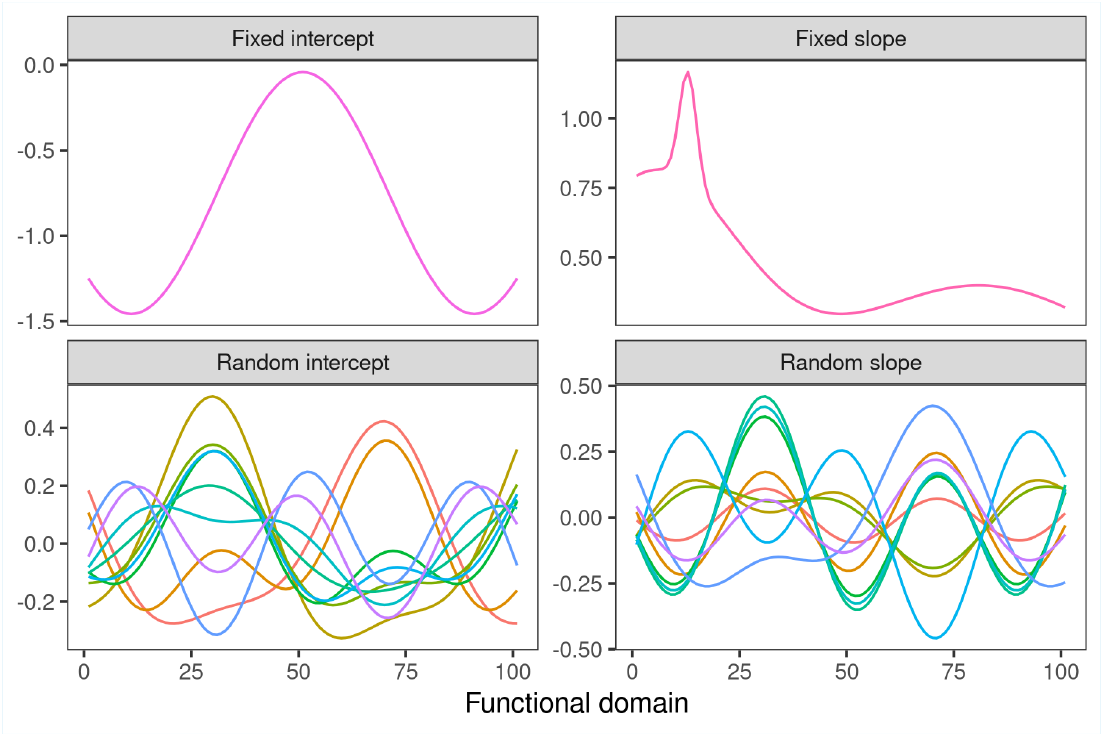
Functions corresponding to the fixed effects and a sample of 10 i.i.d. draws of subject-specific random effects. The fixed intercept *β*_0_(*s*) is the true relationship between signal and time when velocity is zero at time *s*, and the fixed slope *β*_1_(*s*) is the true relationship between signal and velocity at time *s*. The random intercept *γ*_*i*,0_(*s*) is individual perturbation of the fixed intercept for mouse *i* at time *s*, and the random slope *γ*_*i*,1_(*s*) is the individual perturbation of the fixed slope for mouse *i* at time *s*.

To create the data and fit the model shown in Figure 1, we generated *I* = 40 unique mice with *J* = 40 replicates each. For clarity, only 3 unique IDs and 10 replicates are shown in Figure 1-iB. The random effects shown in Figure 1Bii- are synthetic for visualization purposes, but the true random effects and instantaneous covariate recordings are shown below, in Figure 2.

**Appendix 1–figure 2.**
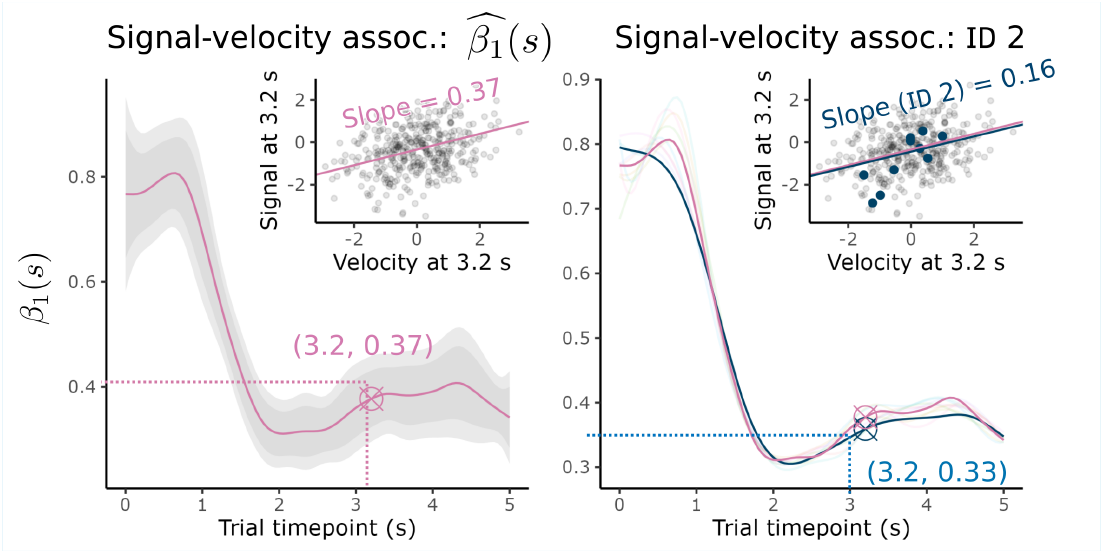
Data points in the insets correspond to the true values generated for the demonstration.

## Appendix 2

### 95% Confidence interval coverage in simulations

We simulated data with the procedure described in Appendix 1. For results in Verification of method performance in simulations, we generated data according to the following mixed effects model:

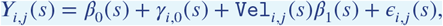

where the functional covariate, fixed effects, and random effects were defined in Appendix 1. We removed the random slope by setting *γ*_*i*,1_(*s*) = 0 for all values of *s*.

### Coverage with random slopes

We also examined coverage for simulated data that included random slope, i.e., with data generated according to the mixed effects model

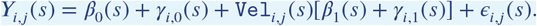

We found that the joint 95% CI coverage of cFLMM models fit to data simulated with random slopes was more sensitive to the magnitude of between-subject variability (i.e., `SNR`_*B*_, the variance of the random effects). For example, in Figure 1, higher variance of the random effects was associated with lower joint coverage. The pointwise coverage tended to be far more robust to this parameter. We present simulations without a random slope in the main body to visualize changes in coverage as a function of `SNR`_*σ*_.

**Appendix 2–figure 1.**
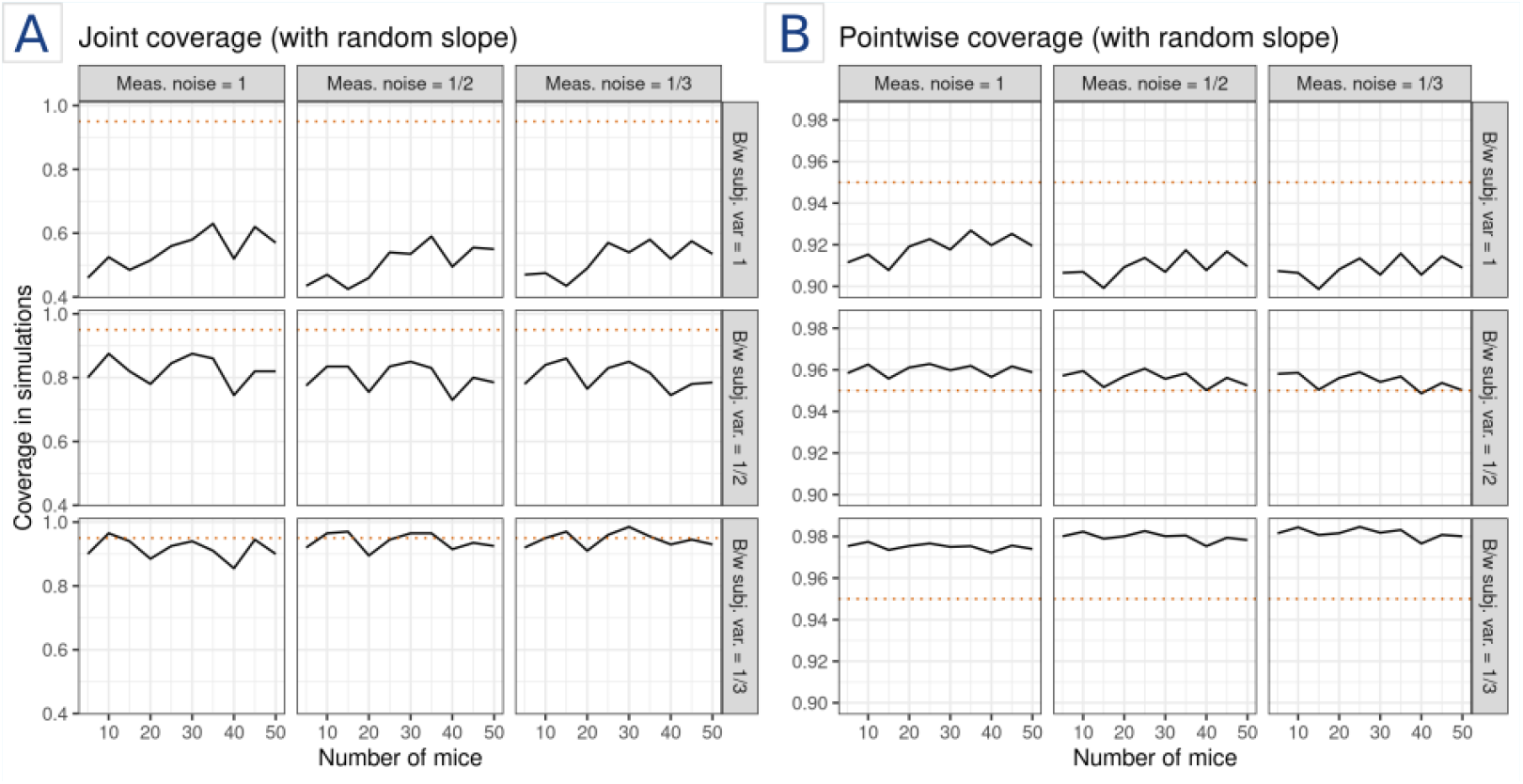
Joint (**A**) and pointwise (**B**) coverage in simulations when incorporating a random slope.

### Visualization of signal-noise ratios

For visual reference, we generated synthetic photometry signals based on the simulation setup (Fig 2).

**Appendix 2–figure 2.**
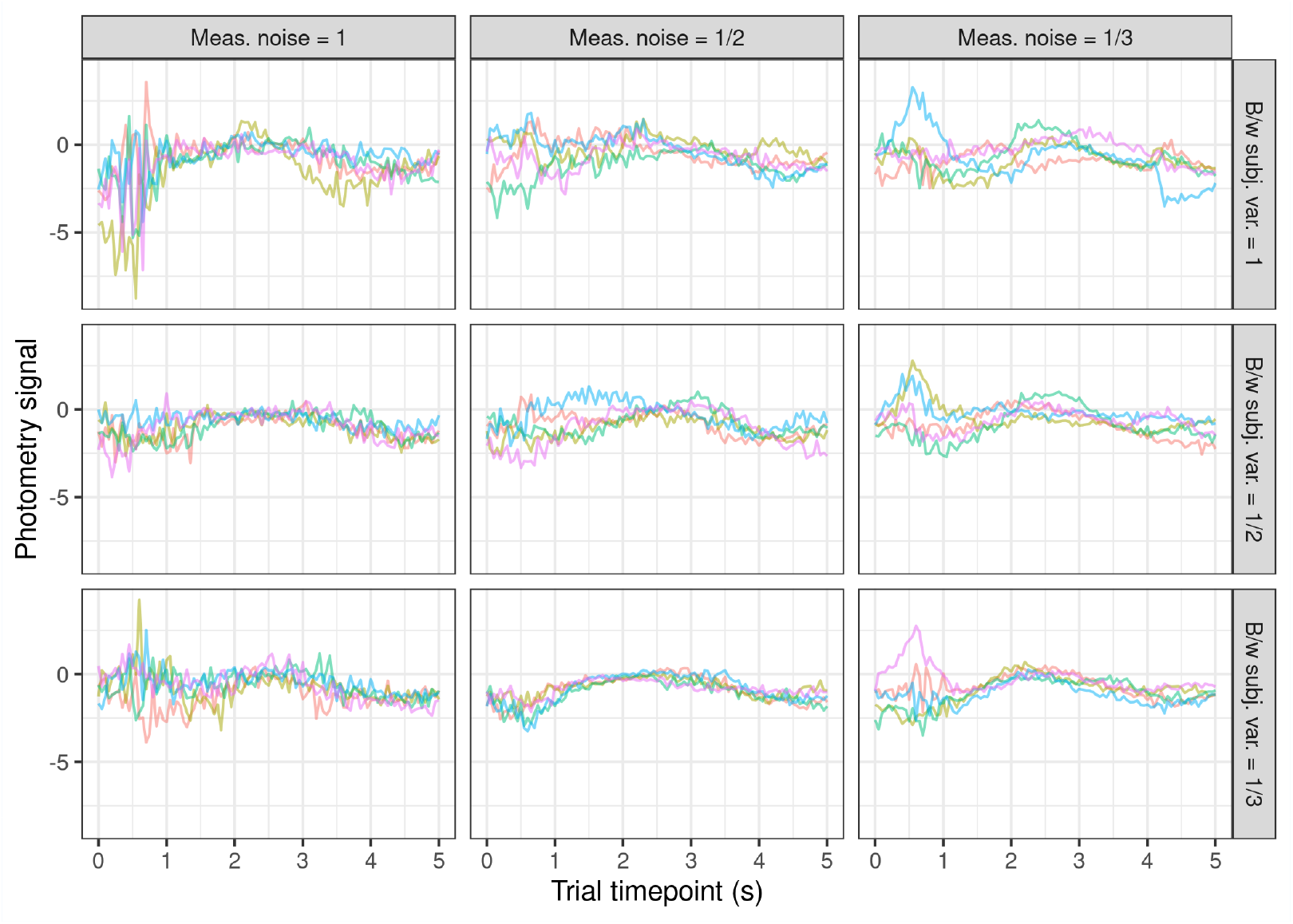
Traces of synthetic photometry signals for five subjects with one observation each.

## Appendix 3

### Additional fixed effects estimates

Although we incorporated non-functional fixed effects for trial and session into the model, we did not present or interpret these estimates in the main body of Modeling behaviors that change over the trial. There are slight differences in coefficient estimates for the fixed effects associated with trial number and session number. We focused on highlighting the particular changes in interpretation associated with exchanging the functional covariate `Lick`_*i,j,l*_(*s*) (which varies over trial timepoints) for the non-functional lick rate summary co-variate, `LR`_*i,j,l*_ (which is a scalar value for each trial).

The magnitudes of the coefficient estimates vary between the cFLMM and ncFLMM models due to the difference in magnitude between the values of `Lick`_*i,j,l*_(*s*), which is binary (0 or 1), and `LR`_*i,j,l*_, which usually takes values within [0, 0.2]. We can, however, still compare the shape and sign (i.e., positive or negative) of the effect estimates. Fixed effect estimates for trial number are consistent across all models, both in the shape of the coefficients and the width of the joint confidence bands. Estimates for session number are also consistent in shape between models, though ncFLMM models for *T* = 0.5, 2.0 have much wider joint confidence bands during early within-trial timepoints compared to cFLMM (Fig 1). This may be related to the use of lick rate as a summary statistic, as increased licking behavior does not immediately follow reward delivery (Fig 2A-i). This inconsistency may introduce additional variance into the coefficient estimates early in the trial. However, it is unclear why this affects confidence band estimates for session and not trial number or lick rate; it may be due to learning having a non-linear relationship with increasing session. We compare the lick covariate functional coefficient estimates in the main text.

**Appendix 3–figure 1.**
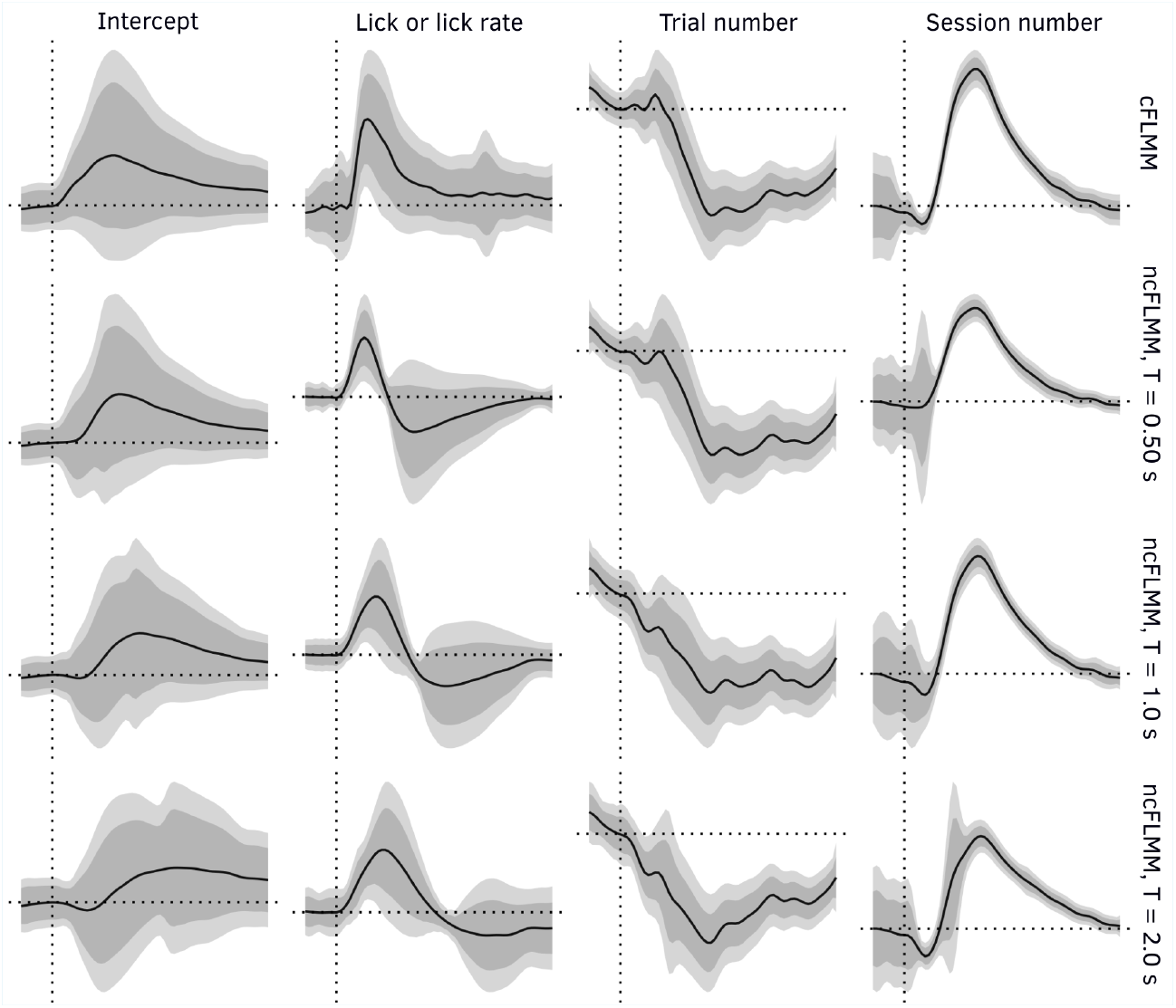
Point estimates and confidence intervals for all fixed effects included in the models compared in Modeling behaviors that change over the trial. Axes are omitted because the scale of the concurrent and non-concurrent models are different based on the covariate of lick or lick rate.

## Appendix 4

### Speeding up covariance calculation with method-of-moments

Discussion of speed-ups will refer to the three-step process of model fitting described in Box 1. When estimating covariance between fixed effects (Eqn. 3), the random effects covariance matrix 𝔾 must be estimated from the data. To do so, we apply the method of moments estimator described in ***Greven et al. (2010***), applied in ***Cui et al. (2022***) for this functional mixed modeling estimation strategy, and extended in ***Loewinger et al. (2025***) to general random effect specifications. Specifically, we calculate

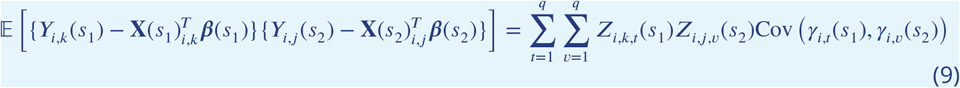

where *t, v* are random-effect covariate indices and *j, k* are trial indices. We can then regress the residual produces onto the random effect covariate products. Compared to the method of moments estimator applied in the non-concurrent FLMM (***Cui et al.,2022***; ***Loewinger et al.,2025***), we substitute **X**(*s*) for **X** and **Z**(*s*) for **Z** in each occurrence.

Although both the non-concurrent and concurrent case theoretically require the same number of calculations, in practice we can reduce the number of necessary calculations in the non-concurrent case as the design matrices are constant across the functional domain. After obtaining the fixed coefficient point estimates 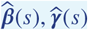 for each point *s*, the univariate models can be discarded in ncFLMM estimation.

In the concurrent case, however, we need to store the functional matrices **Z**(*s*), **X**(*s*) for all *s* for the variance estimation at Step 3. When incorporating even simple indicator variables for categorical random variables, the number of calculations needed to calculate the unique combinations of *s*_1_, *s*_2_ grows quadratically with respect to the size of the functional domain. We can, however, modify the estimator to work with only the row sums of the random effects, using an adaptation of the derivation described in *Appendix 3*.*1* of ***Loewinger et al. (2025***). We briefly review the necessary adaptation for the cFLMM case here.

If *q* is the number of random effects in the model, let *q*^*∗*^ be the number of unique random effect distributions, where *q*^*∗*^ ≤ *q*. For clarity, if we have some *r* ∈ {1, 2, …, *q*^*∗*^} corresponding to a categorical variable with *r*^′^ groups, we have a corresponding set ℐ_*r*_ ⊆ {1, 2, …, *q*} of size *r*^′^ − 1 indexing the columns in the random effect design matrix containing the indicator variables necessary to model *r*. Let *r, m* index the sets of random effects with the same distribution. We can then rewrite Equation 9 as

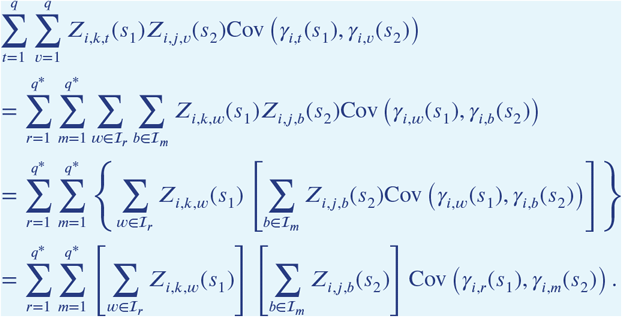

We derived the final equality by noting for random effects in the set ℐ_*r*_, we have 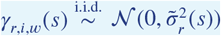, and thus 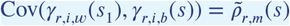, where 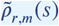 is the same for all *w* ∈ ℐ_*r*_, *b* ∈ ℐ_*m*_. Thus, during Step 1, the matrices **Z**(*s*) can be reduced to a vector of row sums, summed across the vectors **Z**_*i,k*_(*s*), saving both memory and computational time during the covariance estimation step.

Unfortunately, there are no similar simplifications for the **X**(*s*) in Equation 3 that we are aware of, and so these design matrices must be stored in memory until Step 3. Luckily, the matrices *X*(*s*) tend to be far smaller than the *Z*(*s*) and are thus usually manageable to store in memory. Moreover, the need to keep the *X*(*s*) in memory is largely a byproduct of the choice to incorporate the smoothed estimates of coefficients in the variance estimation of Step 3. In future model development, it may be possible to parallelize Steps 1 to 3 across the functional domain and eliminate the need to store all *L* instances of **X**(*s*) simultaneously.

## Footnotes

1 Shifting and scaling does changes the magnitude of estimated coefficients but not the shape. We recommend scaling when covariates have very different magnitudes. E.g., here, trials range from 1 to 100 while licks range from 0 to 1.

## References

Cui E, Leroux A, Smirnova E, Crainiceanu CM. Fast Univariate Inference for Longitudinal Functional Models. Journal of Computational and Graphical Statistics. 2022; 31(1):219–230. 10.1080/10618600.2021.1950006, doi: 10.1080/10618600.2021.1950006, pMID: 35712524.

Greven S, Crainiceanu C, Caffo B, Reich D. Longitudinal functional principal component analysis. Electronic Journal of Statistics. 2010; 4(None):1022–1054. 10.1214/10-EJS575, doi: 10.1214/10-EJS575.

Jeong H, Taylor A, Floeder JR, Lohmann M, Mihalas S, Wu B, Zhou M, Burke DA, Namboodiri VMK. Mesolimbic dopamine release conveys causal associations. Science. 2022; 378(6626):eabq6740. https://www.science.org/doi/abs/10.1126/science.abq6740, doi: 10.1126/science.abq6740.

Loewinger G, Cui E, Lovinger D, Pereira F. A statistical framework for analysis of trial-level temporal dynamics in fiber photometry experiments. eLife. 2025 Mar; 13:RP95802. doi: 10.7554/eLife.95802.

Machen B, Miller SN, Xin A, Lampert C, Assaf L, Tucker J, Pereira F, Loewinger G, Beas S. The encoding of interoceptive-based predictions by the paraventricular nucleus of the thalamus D2+ neurons. bioRxiv: The Preprint Server for Biology. 2025 Mar; p. 2025.03.10.642469. doi: 10.1101/2025.03.10.642469.

Pinheiro J, Bates D, R Core Team. nlme: Linear and Nonlinear Mixed Effects Models; 2025, https://CRAN.R-project.org/package=nlme, r package version 3. 1–168.

Scheipl F, Gertheiss J, Greven S. Generalized functional additive mixed models. Electronic Journal of Statistics. 2016; 10(1):1455–1492. 10.1214/16-EJS1145, doi: 10.1214/16-EJS1145.

